# Charge-State-Dependent Collision-Induced Dissociation Behaviors of RNA Oligonucleotides via High-Resolution Mass Spectrometry

**DOI:** 10.1101/2023.01.29.526146

**Authors:** Rui-Xiang Sun, Mei-Qing Zuo, Ji-Shuai Zhang, Meng-Qiu Dong

## Abstract

Mass spectrometry (MS)-based analysis of RNA oligonucleotides (oligos) plays an increasingly important role in the development of RNA therapeutics and in epitranscriptomic studies. However, MS fragmentation behaviors of RNA oligos are understood insufficiently. In this study, we characterized the negative-ion-mode fragmentation behaviors of 26 synthetic RNA oligos of four to eight nucleotides (nt) in length by collision-induced dissociation (CID) using a high-resolution, accurate-mass instrument. We find that in the CID spectra acquired under the normalized collisional energy of 35%, ∼70% of the total peak intensity belonged to sequencing ions (*a-B, a, b, c, d, w, x, y, z*), ∼25% belonged to precursor ions with either complete or partial loss of a nucleobase in the form of a neutral or an anion, and the remainder were internal ions and anionic nucleobases. Of the sequencing ions, the most abundant species were *y, c, w, a-B*, and *a* ions. The charge state of the RNA precursor ions strongly affected their fragmentation behaviors. As the precursor charge increased from -1 to -5, the fractional intensity of sequencing ions in the CID spectra decreased, whereas the fractional intensity of precursor ions with neutral and/or charged losses of a nucleobase increased. Moreover, RNA oligos containing U, especially at the 3′ terminus, tended to produce precursors that lost HNCO and/or NCO^-^, which presumably corresponded to isocyanic acid and cyanate anion, respectively. These findings build a strong foundation for mechanistic understanding of RNA fragmentation by MS/MS, contributing to future automated identification of RNA oligos from their CID spectra in a more efficient way.

## 1. Introduction

Tandem mass spectrometry (MS/MS) has evolved to become an indispensable approach to elucidate biomolecular structures for both fundamental research in biological processes and widespread applications in biomedicine.^1,2^ Particularly in the last decade MS/MS has been increasingly employed for RNA sequencing and post-transcriptional modification (PTM) characterization.^3-7^ This is primarily due to the high sensitivity, throughput, and mass-accuracy afforded by MS.^3^ In recent years, characterization of RNA oligonucleotides (oligos) by MS/MS has been incrementally adopted during the process to develop oligonucleotide therapeutics.^8-11^ Furthermore, the emerging field of epitranscriptomics, which focuses on the chemical modifications of RNA molecules at a global scale, has also prompted methodological developments in MS technologies.^12-17^ MS is regarded as one of the most reliable techniques to enable exploring PTMs on RNA molecules in a more efficient manner, allowing simultaneous detection of all PTMs on any modified nucleoside.^3^

There have been far fewer previous MS-based investigations of nucleic acids (NAs), compared to those of peptides and proteins, reflecting a long-standing dearth of attention to this area. This is in part due to two main challenges. One is the structural differences between proteins and NAs; the phosphodiester backbone of NAs makes MS analysis better in the negative rather than positive ion mode. However, the encompassing liquid chromatography, MS fragmentation and software package for negative-ion NAs are, so far, far from being routine.^18-21^ The other challenge is that MS dissociation of deprotonated NAs is much more complicated than that of protonated peptides, leading to insufficient understanding of their fragmentation behaviors. This results in technical difficulties and suboptimal performance of the currently available software packages.^20,21^ To the best of our knowledge, many mechanistic studies have been performed that propose models to understand peptide dissociation.^22^ However, only a handful of investigations have focused on short NA fragmentation mechanisms using high-resolution MS.^23-25^ These two limitations have hampered the widespread applications of modern MS to efficient RNA sequencing in a high-throughput and automatic way.^25^

MS studies of monomeric NAs dated back to the early 1960s. Such studies employed electron impact ionization to analyze sample constituents. ^26,27^ In the late 1980s, the advent of soft ionization, electrospray ionization (ESI),^28^ and matrix-assisted laser desorption/ionization (MALDI)^29^ accelerated the pace of developments in MS for oligonucleotide analysis. Most of the early investigations concentrated on the dissociation mechanisms of oligonucleotides that contained one to three nucleotides (nt). Many fragmentation pathways were discovered from analyses of DNA and RNA, including a variety of sequence-determining ions from backbone cleavage, neutral or charged nucleobase losses from precursor ions, and released charged-nucleotides,^30-38^ but these published discrepant data remains a bit turbid for oligonucleotide fragmentation.^30^ Additionally, far fewer mechanistic studies have addressed RNA dissociation than DNA dissociation.^31^

Collectively, previous studies have revealed that many factors can markedly affect oligonucleotide fragmentation behaviors in a mass spectrometer. These factors include both intrinsic features, such as sequence, length, charge state, and terminal group, and extrinsic features, such as the fragmentation technique used and the energy deposited.^23-25^ These studies used short model NAs or analogues under low-resolution instruments. Thus, there is no comprehensive understanding of the fragmentation patterns arising from longer RNA oligos via modern high-resolution MS.

To remedy these gaps in knowledge and establish a fragmentation pattern landscape for MS oligoribonucleotide analysis, we here systematically investigated the unique CID fragmentation patterns from 26 synthetic RNA oligos (containing four to eight nt each) using modern high-resolution MS. Compared with the CID spectra of tryptic peptides, those of RNA oligos were generally much more complicated, more types of fragment ions and a variety of nucleobase losses observed. One of the most significant patterns in oligonucleotide CID was the high dependence on the precursor charge state. For example, there was a pronounced difference in CID spectra between the lowly-charged and the highly-charged precursor ions, even from the same RNA oligo (Figure 2). We acquired a large range of CID spectra from different sequences, each with multiple charge states under several energy conditions. Using the CID spectra from 26 synthetic oligoribonucleotides, we aimed to determine the charge-state-dependent fragmentation behaviors in negative ion mode.

## 2. Experimental Section

### 2.1 Materials and sample preparation

Twenty-six oligoribonucleotides containing four to eight nt each (Supplemental Table S1) were synthesized by the Biological Resource Center of the National Institute of Biological Sciences or Tianjin Wuzhou Kangjian Biotehnology Co., LTD, Beijing. Each sample was dissolved in water to 1 mM, then diluted to 10^−2^ mM in 7.5 mM hexylamine (Energy Chemical, Shanghai, China) with 50% acetonitrile (ACN).^18^

### 2.2 Mass spectrometric analysis

All MS experiments were performed on an Orbitrap Fusion™ Lumos™ Tribrid™ Mass Spectrometer (Thermo Fisher Scientific). Sample solutions were introduced into the electrospray source at a flow rate of 2 μl/min via a needle using a syringe pump. The ion spray voltage was -2.5 kV. The temperature of the ion transfer tube was 320°C. Microscan was set to 3 for both MS1 and MS2. The AGC target was 1E6 for MS1 and 1E5 for MS2. Maximum injection time was adjusted automatically for both MS1 and MS2. The resolution was set to 120k for both MS1 and MS2 spectra. The RF lens was set to 45. MS1 and MS2 spectra were in profile and centroid form, respectively. An inclusion containing all of the theoretical precursor *m/z* values for each RNA oligo from charge -1 to charge -9 was used for each sample. The fragmentation method was CID and the normalized collisional energy (NCE) was varied from 10 to 50, increments of 5.

### 2.3 Data analysis

A custom script was developed in MATLAB to analyze all generated MS data. Theoretical *m/z* values were calculated to the fourth decimal place for the following ions: precursors with all potential nucleobase losses or water loss; internal ions induced by double fragmentation; sequencing ions of *a-B, a-, b-, c-, d-, w-. x-, y-*, and *z*-ions, each with multiple charge states; and precursors with a released base or lost cyanate. All computed *m/z* values were compared with the observed peaks from each CID spectrum. Each observed peak was matched to possible ions; the mass was calculated based on the oligomer sequence with a mass tolerance of 3 ppm. The statistical results of sequencing ions and precursors with lost bases (with respect to the number and relative intensities) were analyzed under each charge state observed.

## 3. Results and Discussion

Many factors can affect the fragmentation behaviors of RNA oligos.^31^ Among them, the precursor ion charge state was the most influential. Using RNA fragment ion nomenclature in Figure 1,^32^ we here show a typical example from our data set (Figure 2). Four charge states from -1 to -4 were observed for this four-nt oligo, 5′-OH-GUCA-OH-3′.

**Figure 1.**
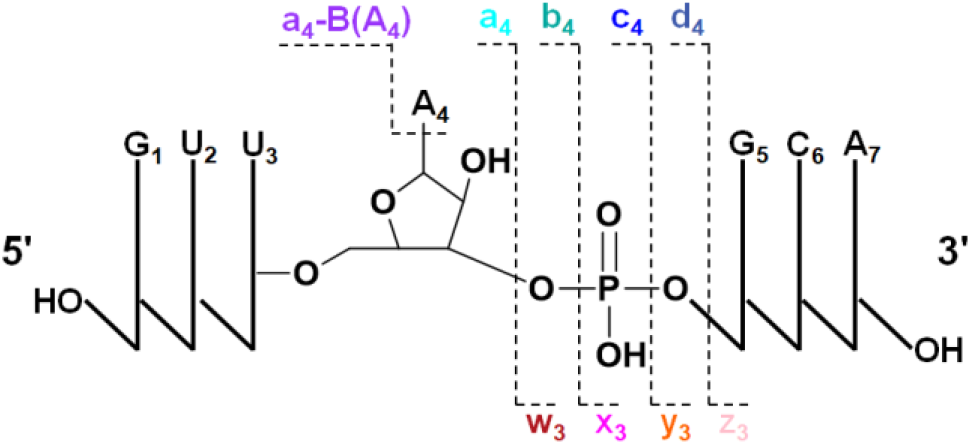
Nomenclature of RNA collision-induced dissociation (CID) fragmentation The sequence of the RNA oligo is GUUAGCA. Both the 5′ and 3′ termini are –OH groups. Ions containing 5′-termini are named *a, b, c*, and *d*. Ions containing 3′-termini are designated *w, x, y*, and *z*. The subscript number after each nucleotide letter indicates its position in the sequence. The subscript number after each ion is the number of nucleotides it contains. ‘B’ indicates the nucleobase released from the corresponding nucleotide after glycosidic bond cleavage.

**Figure 2.**
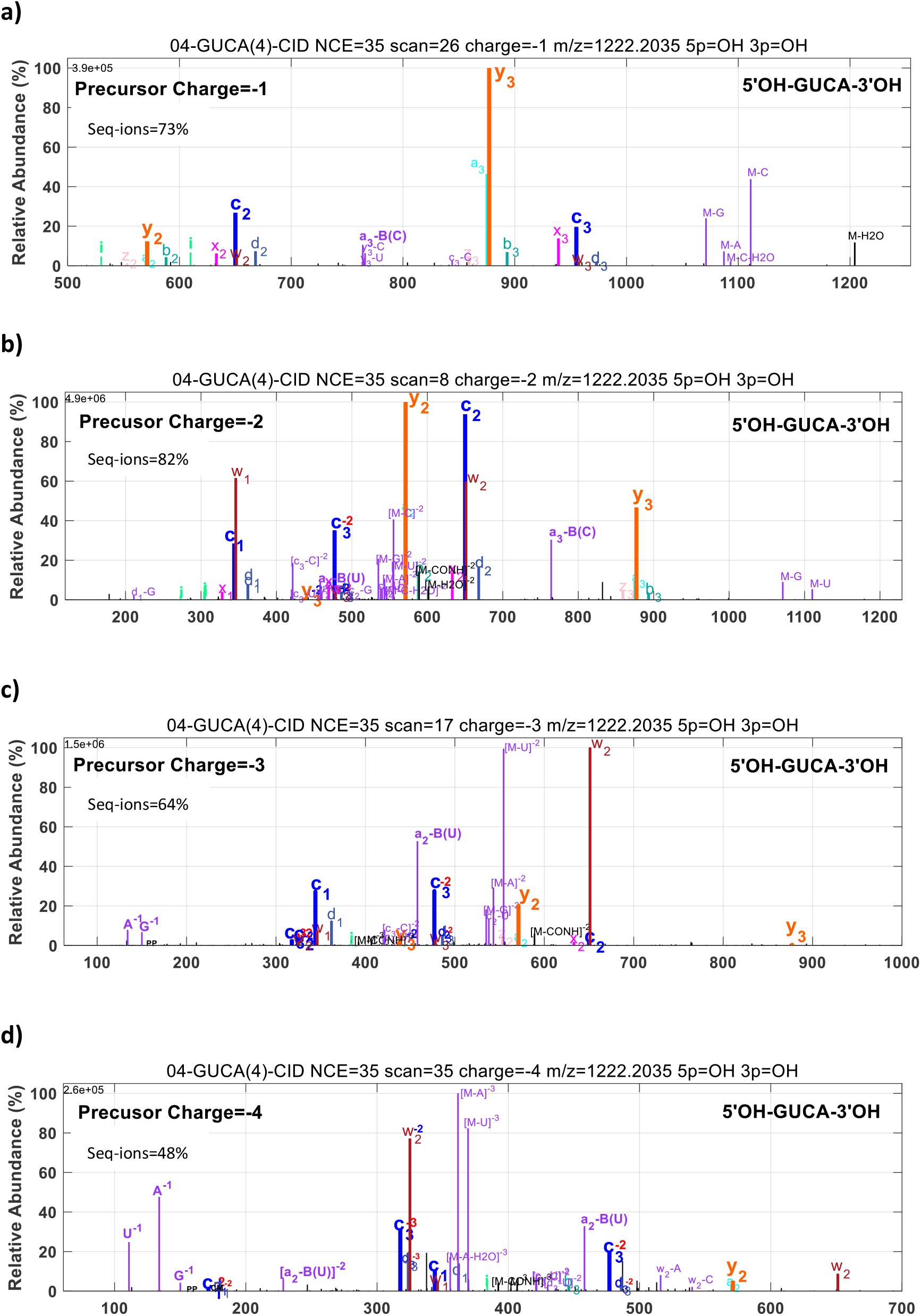
Collision-induced dissociation (CID) spectra from the RNA oligo GUCA, which had – OH groups on both the 5′ and 3′ termini. The charge-state-dependent dissociation was under a normalized collisional energy (NCE) of 35%. Spectra are shown for precursor charges of a) -1, b) -2, c) -3, and d) -4.

As the precursor charge state increased from -1 to -4, the total intensity fraction of all seq-ions first increased in the CID spectra from 73% (−1) to 82% (−2), then decreased to 64% (−3) and 48% (−4) (Figure 2). Specifically, the abundance of *w* ions increased from being undetectable in the -1 spectrum to being prominent in the -4 spectrum. Such precursor-charge-state dependent changes were typical of the RNA oligos examined in this study, as discussed below.

### 3.1 RNA oligos were multiply-charged and their abundance was dependent on the precursor charge-state in negative ion mode

The negative ion mode MS analysis of 26 RNA oligos of four to eight nt in length showed that all these oligos were multiply charged (Figure 3a). There were at least three charge states observed from -1 to -6 for each RNA oligo. The precursor ion abundance was charge-state dependent. Of all charges detected, -2 was the dominant precursor charge state for all 26 RNA oligos. The highest charge observed for each sequence was generally associated with its length; precursors at -3, -4, -5, and -6 were detected for oligos of four, five, six to seven, and eight nt in length, respectively. As the precursor ions tended towards higher charge states along with the increased RNA oligo length, the proportion of -1 precursors among all observed charges decreased (Figure 3b). Concomitantly, the proportion of multiply charged precursors (including all -3/-4/-5/-6 species) increased as the RNA oligo length increased (Figure 3c), although there was relatively high variation when the oligo length was four nt, due to that more -3 precursors were detected for four-nt oligos than for longer ones.

**Figure 3.**
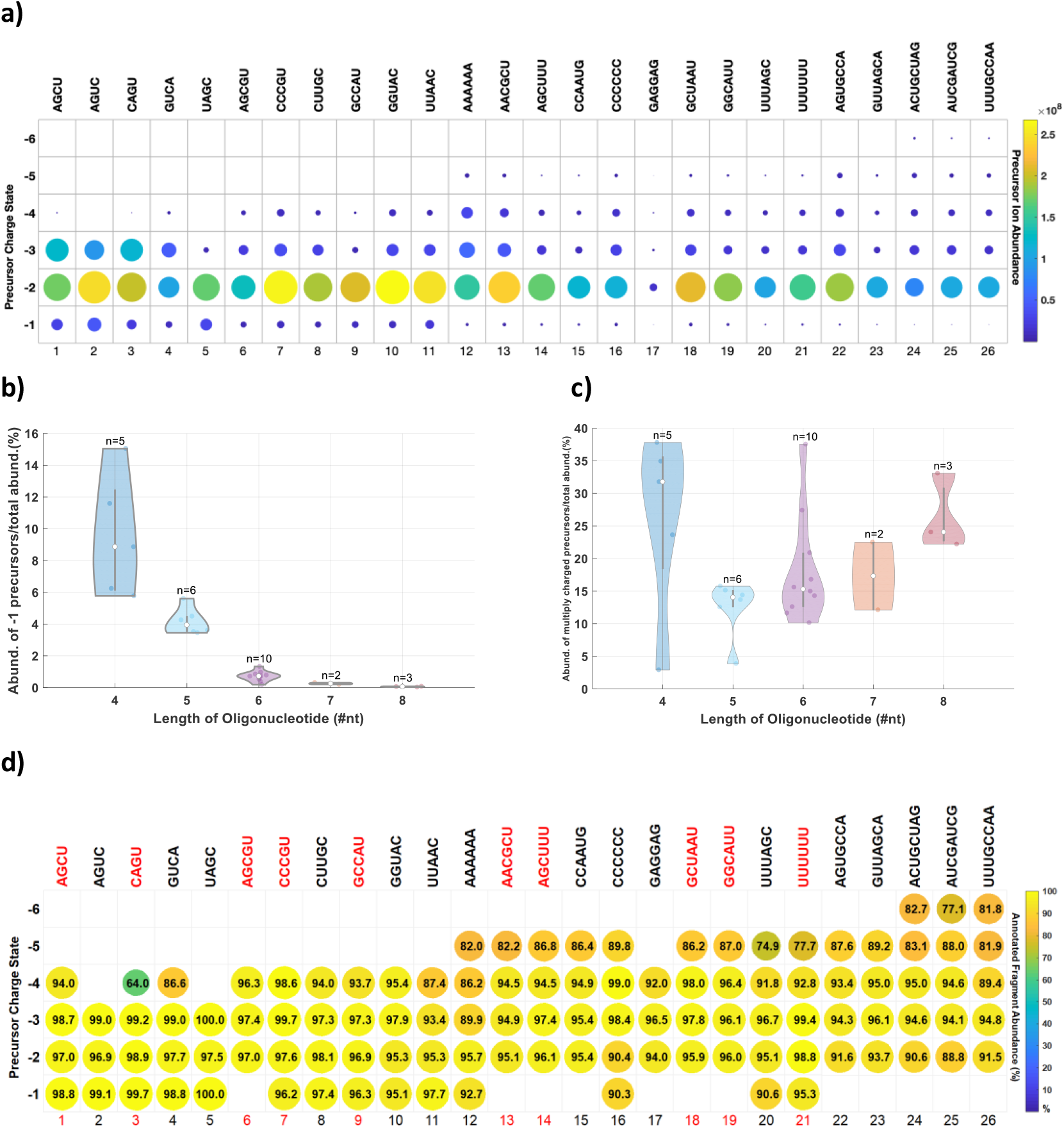
Distribution of precursor ion abundance and annotated fragment fractions of 26 synthetic RNA oligonucleotides (oligos) of four to eight nucleotides (nt) in length under multiple charge states. a) Precursor ion abundance for each of the 26 oligos tested, b) Proportion of -1 precursors corresponding to oligos of each length, c) Proportion of multiply-charged precursors, including all -3/-4/-5/-6 ions observed for oligos of each length. d) Annotated fragment fractions for all 107 collision-induced dissociation (CID) spectra (normalized collisional energy [NCE]=35%). The number *n* on each violin plot in b) and c) indicates the sample number of oligonucleotides tested of that length.

To systematically analyze the ion types and the abundance distributions for all CID fragments, we classified main fragment ions into four categories: sequencing ions (seq-ions), which included nine sub-types (*a-B, a-, b-, c-, d-, w-, x-, y-*, and *z*-ions); nucleobase-lost precursor ions, including isocyanate acid or cyanate anion losses (M-Base/NCO^-^); internal ions (double cleavages); and others, which included released nucleobases and their variants (charged). After automatically annotating each ion from 107 high-resolution CID spectra based on this classification system, we measured the interpretability of each spectrum using the summed abundance of all annotated peaks divided by the summed abundance of all peaks with relative abundance (RA) > 1% (Figure 3d). For lowly- and moderately-charged precursor ions (charge/length ≤ 60%), we achieved an interpretability of > 95% of all fragment ions, producing a highly annotated dataset for further statistical analysis. For highly-charged precursors (charge/length > 60%), the average interpretability was ∼87%, slightly lower than that of the lowly/moderately-charged spectra due to some additional peaks unannotated, such as precursors with double-base losses. Taken together, this dataset showed that CID spectra from RNA oligos, especially from lowly- or moderately-charged precursors could be almost comprehensively explained. Such spectra are therefore highly predictable and have the full potential to be interpreted automatically.

To ensure the high quality of each spectrum, we first acquired CID spectra under NCEs of 10-50% (in 5% increments) for each of the 26 RNA oligos at each charge state detected. We then compared those CID spectra under different NCEs for each RNA oligo. When NCE was 35%, nearly all precursors were fragmented and higher NCEs did not result in significant changes in CID patterns. We therefore selected one CID spectrum at NCE 35% as the representative sample for each RNA oligo under each charge state observed, total 107 spectra for further fragmentation pattern analysis.

### 3.2 Sequencing ion number and abundance were highly dependent on the precursor charge state

For all fragments annotated using the four-class system described above, we calculated the total abundance of each class and the proportion of that class out of the total number of fragments (Figure 4a, left). The seq-ion, M-Base/NCO^-^, and internal ions comprised ∼70%, 25% and 5%, respectively. Seq-ion and M-Base/NCO^-^ were of the greatest interest to us, because the number and abundance of seq-ions determines the performance of RNA sequencing by MS/MS, and the abundance of M-Base/NCO^-^ has crucial influence on the sensitivity and accuracy of RNA sequencing. We therefore focused on the charge-dependence nature of these two types of ions in terms of their abundance distributions.

**Figure 4.**
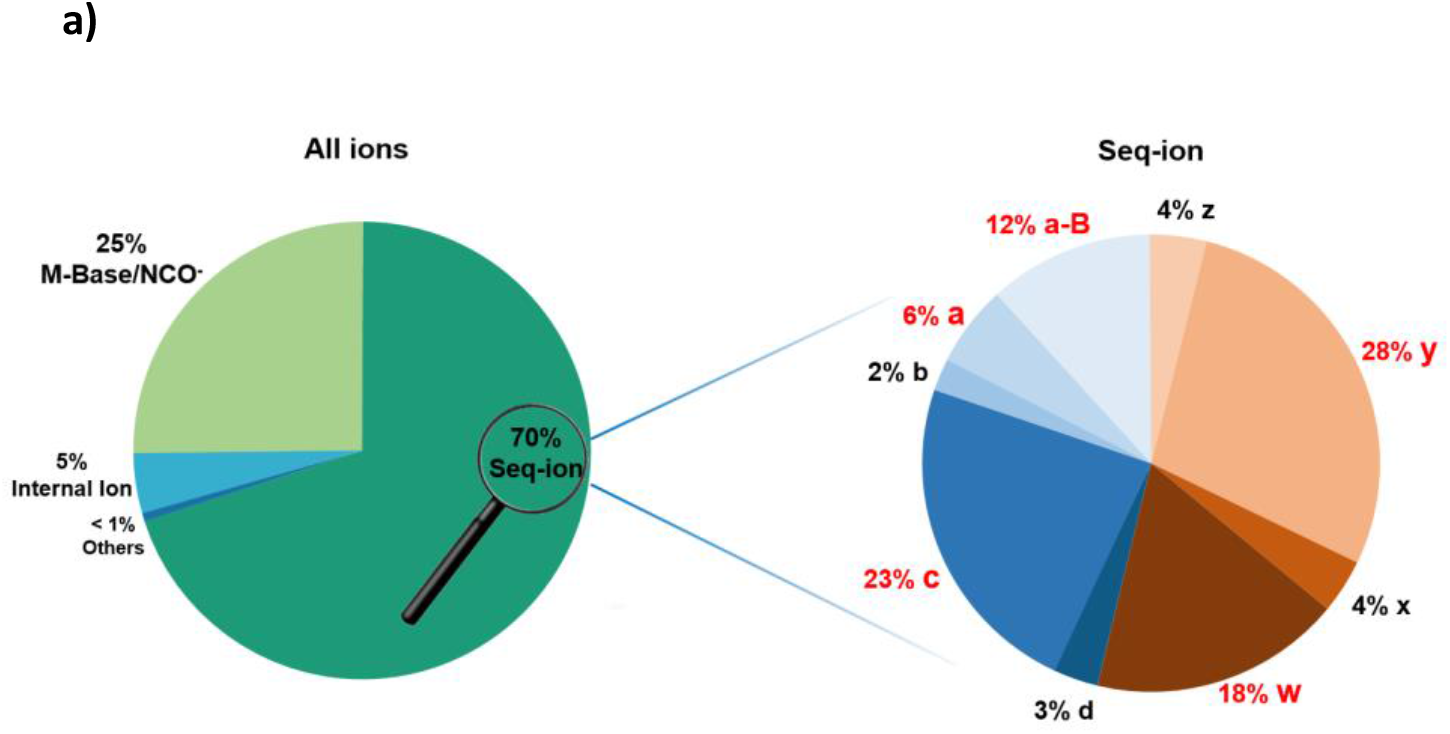
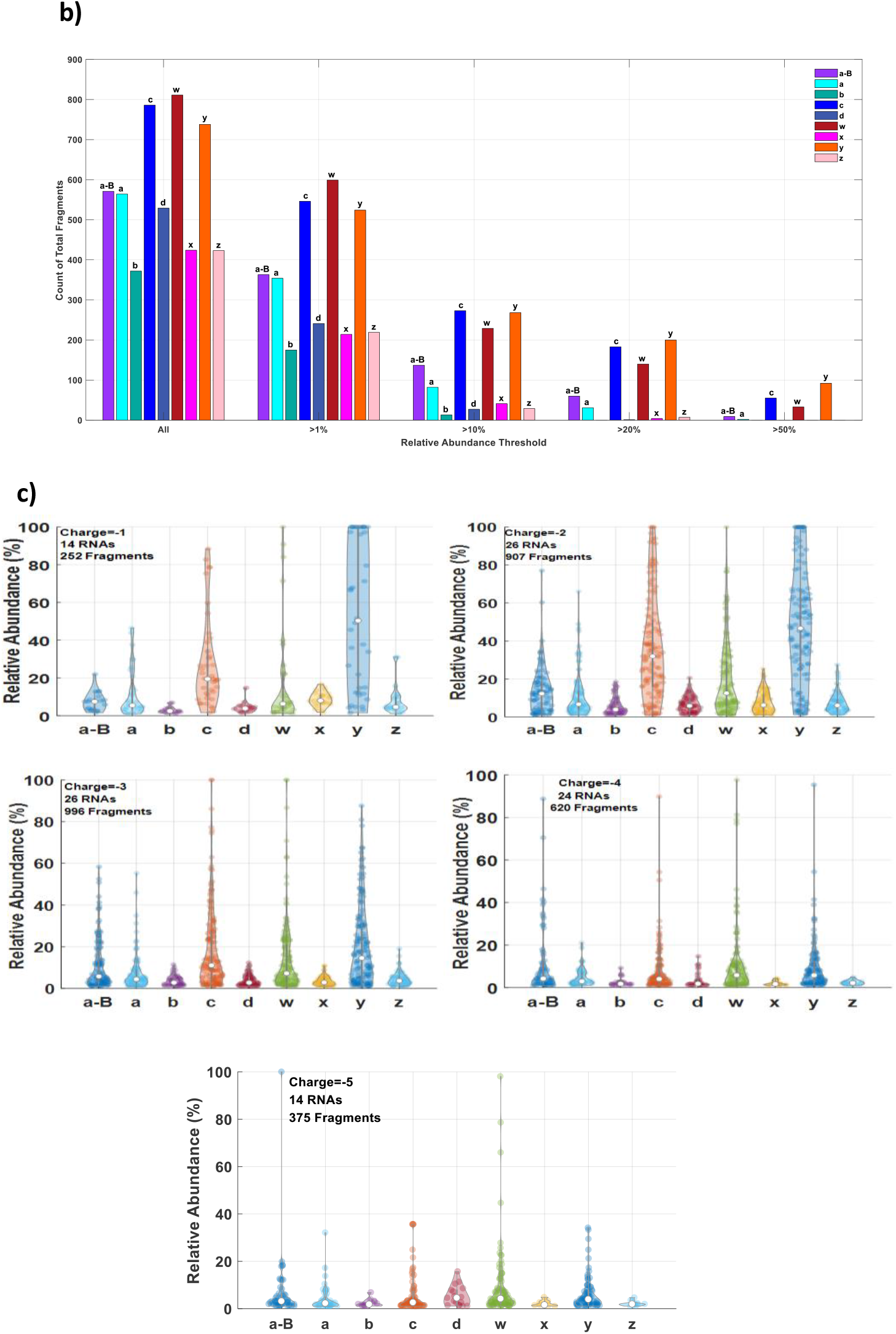
Statistical analysis of fragment number, abundance, and distribution associated with different precursor charge states. a) Proportions of sequencing (seq)-ions, base-lost precursors, and internal ions detected; b) Counts of seq-ions with a relative abundance (RA) greater than 1%, 10%, 20%, or 50%, and when all ions were analyzed, respectively c) Abundance of seq-ions associated with precursors at different charge states.

We next analyzed the total abundance of each sub-type of seq-ion (Figure 4a, right). The five most abundant ions with their abundance coverage were *y* (28%), *c* (23%), *w* (18%), *a-B* (12%) and *a* (6%). Together, these totaled 87% of all seq-ions, and would therefore be among the most critical in an RNA oligo high-throughput search engine. An analysis was then conducted to count the number of each sub-type above an RA threshold (1%, 10%, 20%, or 50%) and all measured ions (Figure 4b). We found that *y, c, w, a-B*, and *a* were the most highly abundant, consistent with the results of the previous abundance-based analysis (Figure 4a). Highlighting the relative abundance distribution of each seq-ion (separated by precursor charge state), could reveal the effect of charge state on the abundance distribution.

We then analyzed the relative abundance of nine types of seq-ions under five charge states from -1 to -5 (Figure 4c). For lowly- and moderately-charged precursor ions (−1 to -3), the median RA of *y*- and *c*-ions was markedly higher than it was for other types of ions. However, as the charge state increased to -4 or -5, these ions became gradually less abundant. Taken together, these data verified that low precursor ion charge states, such as -1/-2, were the optimal choice among all charge states detected; the corresponding CID spectra from lowly-charged RNA oligos (four to eight nt) provided a higher abundance of seq-ions, particularly *y* and *c* ions, which reveal the key information for accurate RNA sequencing.

Of the nine sub-types of seq-ions, RA of *w* ions was more relevant to the nucleotide identity around the cleavage site and the precursor charge state (Figure 5). When the precursor charge was low (−1 or -2) and the 5′ side of the cleavage site was a cytosine, *w* ions were much more abundant than when other combinations of nucleotides were present around the cleavage site (Figure 5a). This trend was diminished when the precursor charge state was high. The discovery of this pattern will help to improve specificity in matching CID spectra to RNA sequences, and prompt the corresponding chemical mechanism exploration.

**Figure 5.**
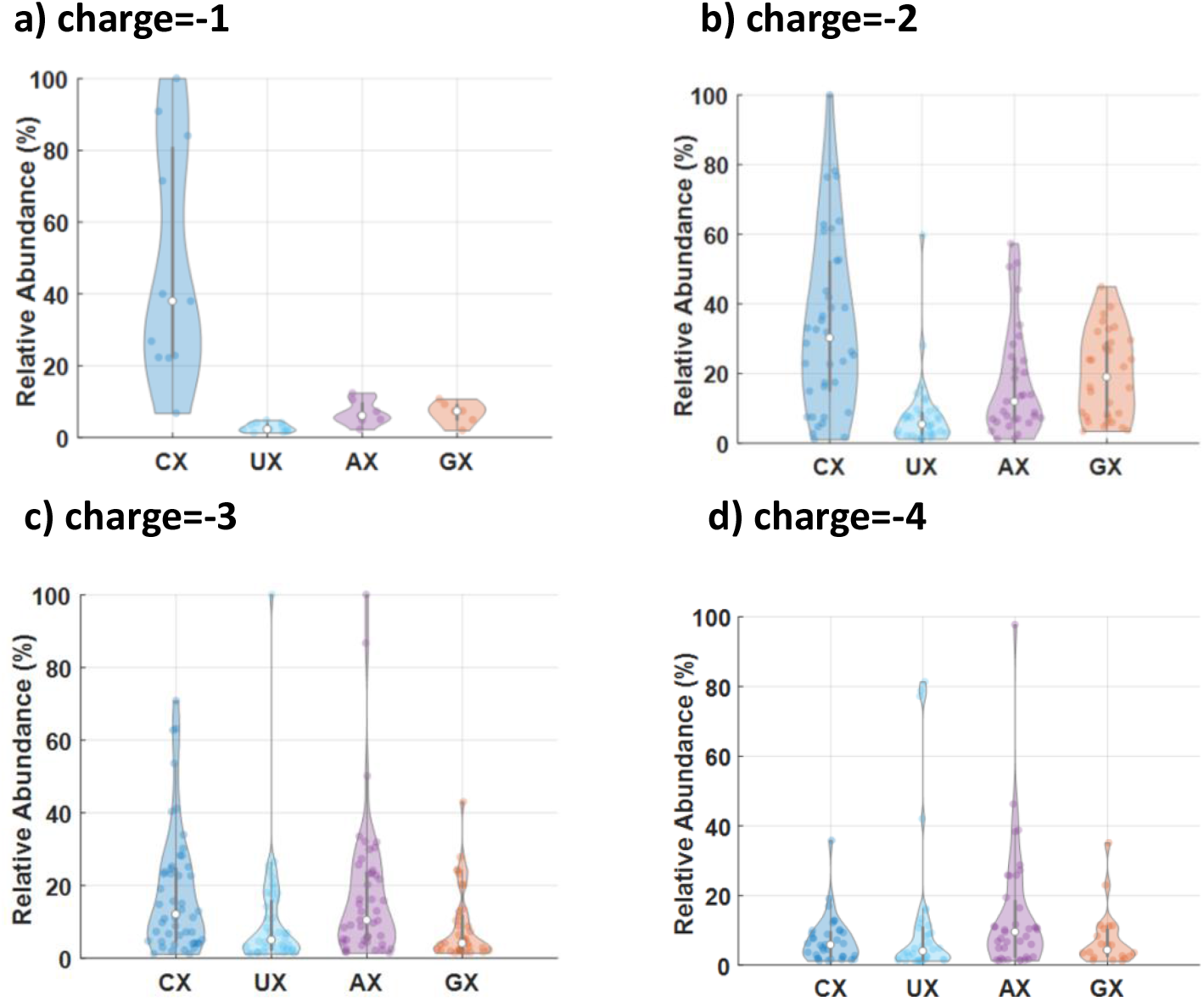
*w* ions were highly abundant when the 5′ cleavage site was a cytosine, and the precursor charge was low (−1 or -2). Here, X stands for any base (C, U, A, or G). Data are shown for fragments with their precursor charge of (a) -1, (b) -2, (c) -3, (d) -4.

### 3.3 Nucleobase losses from precursors were markedly increased at high precursor charge states

Unlike side-chain losses from amino acids in peptides or proteins, nucleobase losses from precursor ions were frequently observed in the CID spectra of RNA oligos, particularly from highly-charged precursors. This phenomenon can be primarily attributed to increases in Coulomb repulsion as a result of more nucleobase charges, which promotes base release via glycosidic bond cleavage.^39^ Uninformative base losses have a harmful impact on the performance of RNA sequencing by MS; base losses can reduce the relative signal-to-noise ratio of seq-ions, which are the most informative in RNA sequencing by MS/MS.

To systematically characterize the charge-state-dependent behaviors of both neutral and charged nucleobase losses, we first carried out a statistical analysis to identify the base peak, i.e., the most abundant peak for each CID spectrum (Figure 6a). For this analysis, we used one representative CID spectrum for each RNA oligo under each charge state, selecting a total of 107 CID spectra corresponding to the 26 RNA oligos.

**Figure 6.**
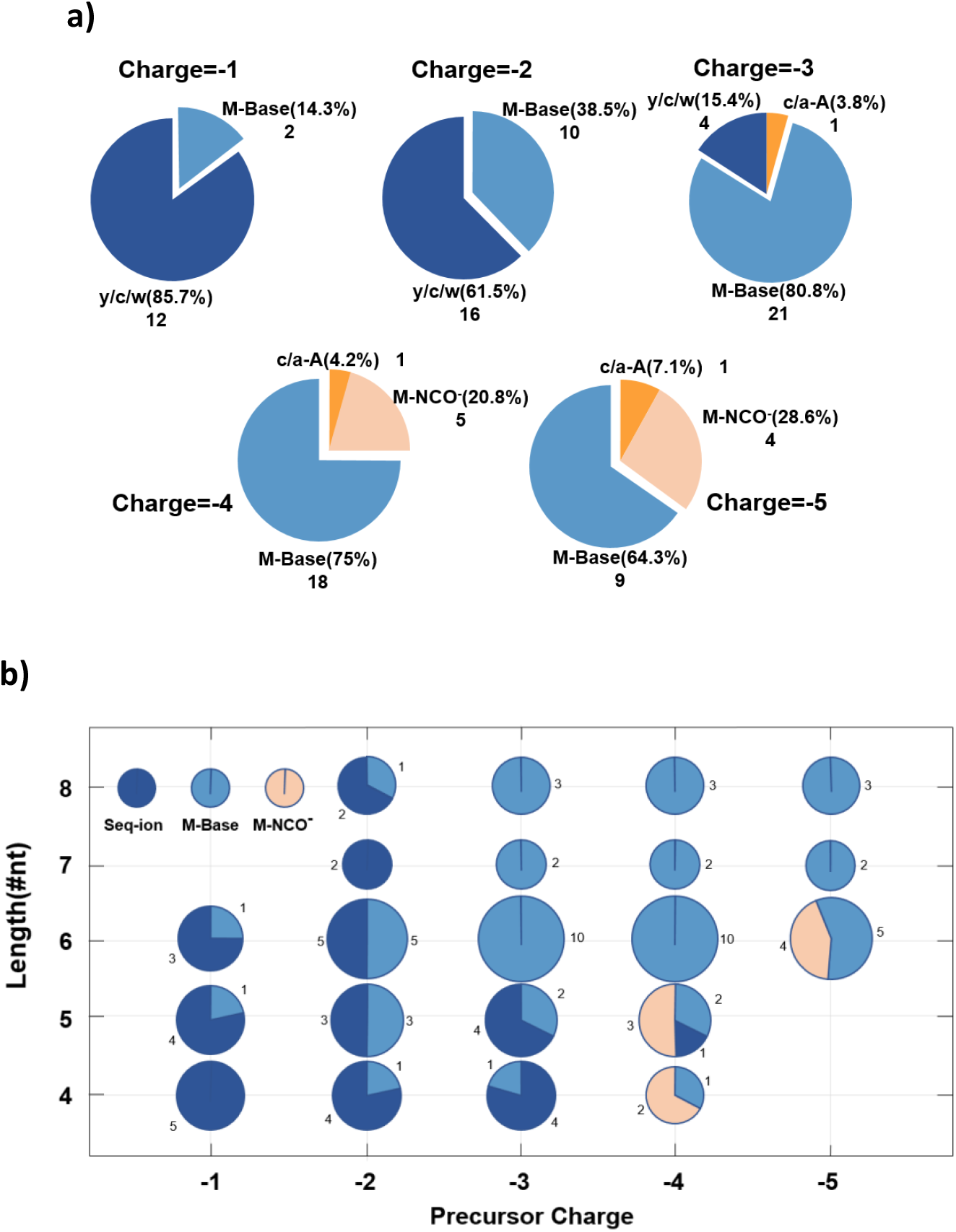
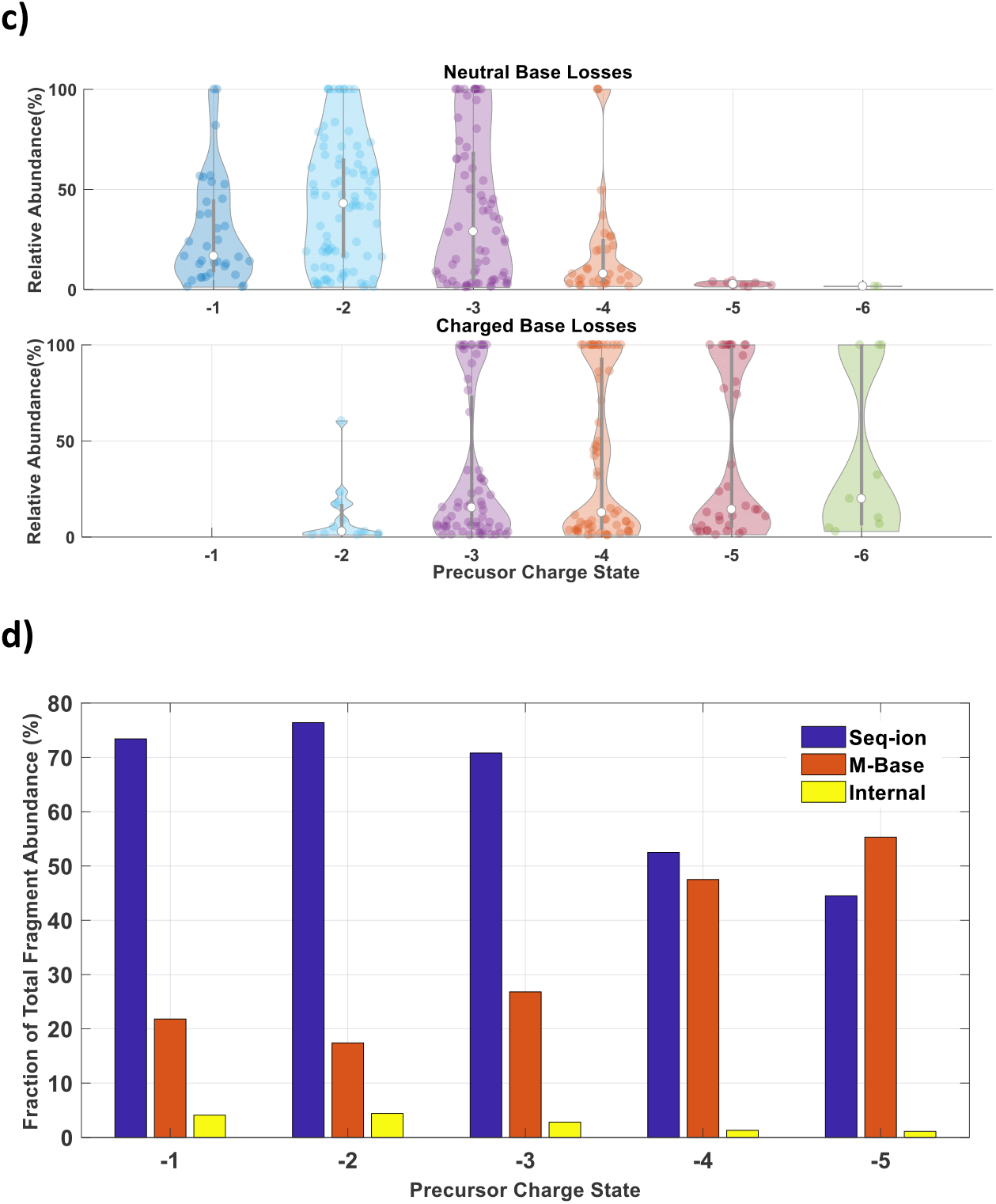
Neutral and charged nucleobase losses from precursor ions. a) Statistics for the identity of the base peak at each charge state. Seq-ions, such as *y/c/w* ions, are shown in deep blue; nucleobase-lost precursors (M-Base) are shown in light blue; cyanate anion-lost precursors (M-NCO^-^) are shown in light orange; and nucleobase-lost seq-ions such as *c/a*-A are shown in dark orange. b) Statistics for the identity of the base peak at each charge state and RNA oligonucleotide length (from four to eight nucleotides). c) Relative abundance (RA) of neutral and charged base losses from precursors at each charge state (−1 to -6). d) RA percentage of all seq-ions, M-Base and internal ions at each precursor charge state. Each integer in a) and b) is the number of the CID spectra whose base peak belongs to that type specified. The corresponding percentage was included in the parenthesis for figure a).

There were four types for the identity of a base peak: seq-ions, such as *y/c/w* ions; nucleobase-lost precursors (M-Base); cyanate anion-lost precursors (M-NCO^-^); and nucleobase-lost seq-ions, such as *c/a*-A. We also analyzed the data by considering the length of each RNA oligo (Figure 6b). These analyses revealed a decrease in the proportion of seq-ions, whereas the proportion of nucleobase-lost precursors increased when the precursor ion charge changed from low to moderate and from moderate to high. As RNA oligo length increased from four, five, or six nt to seven or eight nt, cyanate-lost precursors as the base peak in our data set were not observed. This was likely because few RNA oligos in our dataset contained seven or eight nt, the sequences of those that were present did not have the characteristics that we found to increase the likelihood of producing cyanate anion loss.

We also observed the coexistence of both neutral and charged nucleobase losses in most CID spectra regardless of their abundance. Comparisons between these two types of losses indicate that neutral base losses occurred much more frequently than charged base losses under low charge states, such as -1 or -2 (Figure 6c). However, under high charge states, such as -5 or -6, charged base losses became dominant. For moderate charge states, such as -3 or -4, neutral and charged base losses were similar in their abundance. The phenomenon of precursor charge-state-dependent base losses revealed that highly-charged RNA oligos increased the Coulomb repulsion between a charged nucleobase and the sugar, thus improving the chance of nucleobase release via glycosidic bond cleavage.^39^

We then integrated seq-ion and nucleobase loss behavior data to determine how these trends changed quantitatively with the charge state. The data showed that the fractional abundance of seq-ions in the CID spectra decreased from 73% to 45%, whereas the neutral and charged nucleobase-loss precursor ions increased from 22% to 55%, when the charge state increased from -1 to -5 (Figure 6d). These findings further verified that the lowly-/moderately-charged RNA oligos were the optimal choice for generating CIDs in MS/MS sequencing.

### 3.4 Cyanate anion losses from RNA oligos with 3′-U were dominant under highly-charged states

In addition to the frequently observed intact nucleobase losses from precursor ions of RNA oligos as summarized previously, we also investigated two kinds of partial base losses, in which the base ring opened, and a group released. The released groups here were presumed to be the isocyanic acid (neutral) and the isomer cyanate anion; these have occasionally been reported in the literature in the context of RNA/DNA oligos’ CID spectra.^39^ However, the dissociation behaviors and the specificity of their occurrences had not been comprehensively reported in a large data set. Moreover, the isotopic distributions between the theoretical and experimental isotopic clusters from cyanate-lost precursors were shown to be inconsistent.^39^ Our data sets expanded on previous results and provided accurate isotopic cluster matches. We compared the occurrence frequency of these two types of partial base losses between different oligo sequences under several charge states.

We analyzed both the similarities and the mass deviations between the experimental and theoretical isotopic distributions for the first three highly abundant peaks of each oligo, using UUUUUU as an example (Figure 7a). For that oligo, the base peak was identified as a cyanate-loss precursor. The experimental isotopic distribution was highly consistent with the theoretical cluster. Based on the 107 CID spectra from the 26 synthetic RNA oligos, we analyzed the relative abundance distributions for all isocyanic acid or cyanate anion losses with each of four different 3′ nucleotides (C, U, A, and G) in the 3′ position (Figure 7b) and compared RA values between isocyanic acid (HNCO) and cyanate losses (NCO^-^) under six different charge states (Figure 7c). From these results, we find that the cyanate anion losses were frequently observed to be highly abundant when two conditions were simultaneously met; the original sequence contained a 3′-U (ten red sequences in Figure 3d) and the precursors were highly charged (e.g., -4/-5) (Figures 7b and 7c).

**Figure 7.**
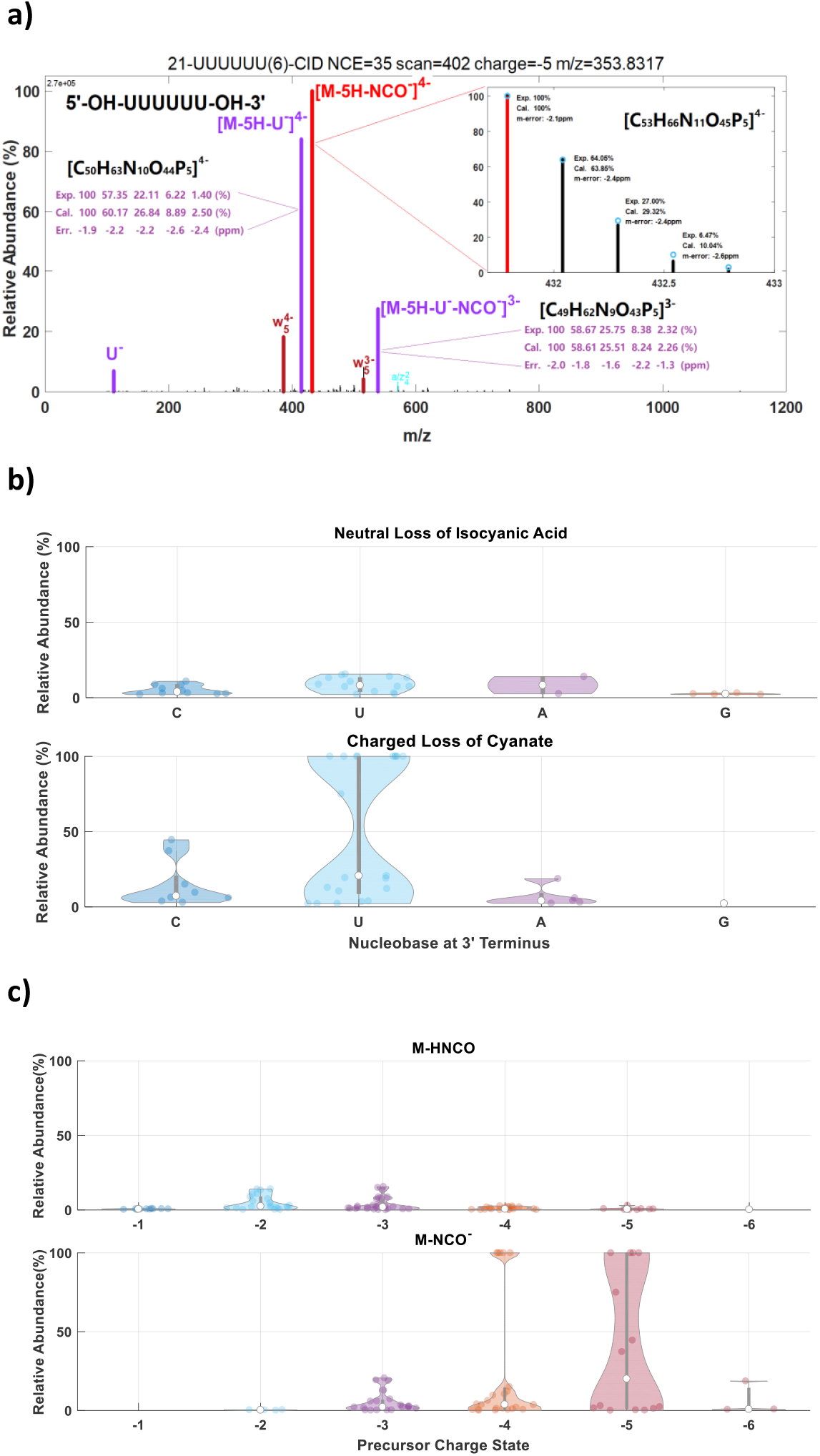
Cyanate anion and isocyanic acid losses from precursor ions a) Example spectrum for the RNA oligonucleotide UUUUUU. Experimental and theoretical isotopic distributions with mass errors are shown for the first three highly abundant peaks. In the zoomed panel, the blue circles correspond to the predicted isotopic distribution. b) Relative Abundance (RA) distributions of precursors with lost isocyanic acid or cyanate anion based on the nucleobase at the 3′ terminus. c) Comparison of RA distributions between isocyanic acid and cyanate anion losses at each precursor charge state.

#### A simplified procedure for assigning isobaric ions

Unlike the distinct-mass nature of N-terminal and C-terminal ion masses observed in peptide fragmentation, the 5′ and 3′ RNA ions are isobaric if the compositions of those ions are the same. For example, the masses of the *a*_*4*_-ion and the *z*_*4*_-ion of the oligo CUAGC are equal if the terminal groups are identical. Therefore, for all palindromic sequences, whose subsequence with equal length is the same whether we read it from 5′ terminus or from 3′ terminus, such as CUAGGAUC, we cannot differentiate between a peak from the 5′ and the 3′ site ion. This complicates spectral interpretation, particularly when the number of all ion types is counted. Nevertheless, this phenomenon has minimal impacts on our previous statistical results. We simplified the processing of counting ions by regarding less frequently observed ions as the more highly abundant isobaric partner if their masses were the same. This simplification is reasonable based on the statistical results from all ion types from the sequences with no isobaric ions.

## 4. Conclusions

Understanding the fragmentation behaviors of RNA oligonucleotides in high-resolution mass spectrometry is the cornerstone to develop new methods for high-throughput, accurate RNA sequencing, PTM characterization, and RNA quantification by MS/MS. We here comprehensively analyzed fragmentation behaviors of 26 synthetic RNA oligos using their representative 107 multiply-charged CID spectra; this encompassed six precursor charge states (from -1 to -6), and oligos with lengths of four to eight nt. Based on a highly annotated data set (over 95% total intensity), our results showed that the fragmentation behaviors of RNA oligos in the negative ion mode were much more complicated than those of peptides or proteins, primarily due to the many possible cleavage pathways. However, we determined that RNA oligo fragmentation by CID was highly predictable and strongly depended on the precursor charge states. This dependence led to three key observations. First, sequencing ions were the dominant ion type under low and medium charge states, which provides valuable information for selection of a charge state to trigger optimal dissociation for high-quality sequencing. Second, the nucleobase losses from precursor ions relied significantly on the precursor charge state; nucleobase losses, particularly charged losses, were most abundant when the RNA oligos were highly charged. Third, the dominance of cyanate anion losses, depended on both the precursor charge state and the identity of the base in the 3′ terminal position. Collectively, these findings regarding RNA dissociation in a high-resolution mass spectrometer deepen our understanding of RNA fragmentation mechanisms. This contributes to a strong foundation that will ultimately allow automated identification of RNA sequences from their CID spectra generated from MS/MS, in a more efficient way.

In addition to the precursor charge state, the terminal groups, fragmentation method used, and PTMs present also affect the fragmentation behaviors of RNA oligos. These other areas are thus the subject of ongoing research in our lab. In the near future, with the rapid development of new MS techniques, RNA sequencing and PTM characterization by MS/MS will become an orthogonal, high-throughput and automated technique.

## Acknowledgments

This study was supported by the National Key Research and Development Program of China (Grant 2020YFF01014505), the Ministry of Science and Technology of China, and Beijing Municipal Science and Technology Commission.

